# Integrated Analysis Revealed Hub Genes in Breast Cancer

**DOI:** 10.1101/414532

**Authors:** Haoxuan Jin, Xiaoyan Huang, Kang Shao, Guibo Li, Jian Wang, Huanming Yang, Yong Hou

## Abstract

The aim of this study was to identify the hub genes in breast cancer and provide further insight into the tumorigenesis and development of breast cancer. To explore the hub genes in breast cancer, we performed an integrated bioinformatics analysis. Two gene expression profiles were downloaded from the GEO database. The differentially expressed genes (DEGs) were identified by using the “limma” package. Then, we performed Gene Ontology (GO) and Kyoto Encyclopedia of Genes and Genomes (KEGG) enrichment analysis to explore the functional annotation and potential pathways of the DEGs. Next, protein–protein interaction (PPI) network analysis and weighted gene coexpression network analysis (WGCNA) were conducted to screen for hub genes. To confirm the reliability of the identified hub genes, we obtained TCGA-BRCA data by using WGCNA to screen for genes that were strongly related to breast cancer. By combining the results from the GEO and TCGA datasets, we finally identified 15 real hub genes in breast cancer. Finally, we performed an overall survival analysis to explore the connection between the expression of hub genes and the overall survival time of breast cancer patients. We found that for all hub genes, higher expression was associated with significantly shorter overall survival times among breast cancer patients.

## 1. Introduction

Breast cancer is one the most severe types of tumor worldwide and is the leading cause of cancer death in women[1]. The breast cancer incidence rate varies widely across regions, with rates ranging from 0.194‰ in East Africa to 0.897‰ in West Europe, and is increasing gradually[2]. There are many risk factors associated with breast cancer, including long-term fertility, the use of hormonal contraception, physical inactivity, and alcohol consumption, but its etiology and pathogenesis are still not definitively understood[3].

With the implementation of some large-cohort human tumor genome projects (for instance, The Cancer Genome Atlas (TCGA) and the International Cancer Genome Consortium (ICGC)), an unprecedented amount of genomic data on tumor samples was generated and became highly important in human cancer research[4]. In addition, many smaller-scale cancer projects led by individual institutions also made great contributions and provided large amounts of valuable data, which were deposited into public databases such as the Gene Expression Omnibus (GEO)[5]. These public cancer data will accelerate the comprehensive understanding of the genetics of cancer, facilitating exploration of the underlying mechanisms of cancer and helping to improve diagnostic methods and preventive strategies.

Due to the limitations of experimental techniques, the development and productive application of microarray and sequencing technology brought cancer research into a new era. These high-throughput techniques have been widely used for global gene expression profiling, which reflect the molecular basis of tumor phenotypes and can be used to classify tumors, discover the pathogenic genes of tumors, explore tumorigenesis and distinguish the occurrence and progression of tumors[6]. A large number of gene microarray datasets in public databases facilitates comprehensive analyses of gene expression in cancer. By using bioinformatics methods and associating the results with clinical data, new biomarkers could be found for the diagnosis, therapy and prognosis of cancer.

In this study, we obtained and used two original microarray gene expression datasets, GSE10810[7] and GSE65194[8], from the GEO database. Strict calibration and filtering were used to obtain differentially expressed genes (DEGs). Then, by a series of bioinformatics approaches, hub genes were identified, and enrichment analysis was used to find possible key pathways of breast cancer. We also downloaded breast cancer gene expression data from TCGA and performed the same strategy to verify our results. A series of Kaplan-Meier (KM) survival plots were also constructed to reveal the connection between hub genes and the prognosis of breast cancer. Finally, we identified 15 true hub genes that closely related to breast cancer. We expect this work to provide further insight into the tumorigenesis and development of breast cancer at the molecular level and provide more precise, practically valuable markers for the diagnosis, therapy, monitoring and prognosis of breast cancer.

## 2. Materials and Methods

### 2.1 Datasets

The GEO is a public database of gene expression profiles and sequence-based data and is freely available for users. Two gene expression profile datasets (GSE10810 and GSE65194) were downloaded from the GEO. Both GSE10810 and GSE65194 were obtained from the GPL570 platform [HG-U133_Plus_2] Affymetrix Human Genome U133 Plus 2.0 Array. GSE10810 includes 58 samples, including 31 breast cancer samples and 27 normal samples, while GSE65194 includes 130 breast cancer samples and 11 normal control samples.

We also downloaded RNA-sequencing gene expression data from TCGA on the UCSC Xena (https://xena.ucsc.edu/) “GDC TCGA Breast Cancer (BRCA)” cohort, which includes 1104 breast cancer samples and 113 normal samples.

### 2.2 Filtering of differentially expressed genes (DEGs)

The “limma” R package[9] was applied to filter the DEGs between the breast cancer patient group and the normal group. The P-value of each DEG was calculated and then adjusted by the Bonferroni method. The threshold that used to DEGs was |log fold change (FC)| ≥ 2 and a Bonferroni P-value <0.01.

### 2.3 Kyoto Encyclopedia of Genes and Genomes (KEGG) pathway and Gene Ontology (GO) enrichment analysis of DEGs

KEGG database is used for systematically analyze and annotate gene functions[10]. The GO database provides a classification of genes into three functional groups: the molecular function (MF) group, biological process (BP) group, and cellular component (CC) group[11]. In this study, KEGG pathway and GO enrichment analyses were conducted by using the “clusterProfiler”[12] R package with a cutoff P-value of 0.05 on the DEGs that we obtained in the previous step.

### 2.4 Integration of protein-protein interaction (PPI) network and cluster analysis

The Search Tool for the Retrieval of Interacting Genes (STRING)[13] is a biological database for predicting pairs of PPIs. We evaluated the interactive relationships of DEGs by STRING and defined the genes with a combined score > 0.9 as key DEGs. Then, we used Cytoscape to develop the PPI network of the key DEGs that we identified. Molecular Complex Detection (MCODE)[14], a plugin for Cytoscape, was used with the default parameters to identify the most important module of the PPI network.

### 2.5 Coexpression network construction and analysis of clinically significant modules

The coexpression network was established by WGCNA, an R package for the construction of weighted gene coexpression networks[15].

In this study, an automatic one-step network construction and module detection method in WGCNA was performed with the default settings, including a correlation type of Pearson, an unsigned type of topological overlap matrix (TOM), a merge cut height of 0.25 and a default minimal module size. The first principal component calculation module eigengene (ME) was used to quantify the coexpression similarity of entire modules. To assess the potential associations between ME and phenotype (case or normal), Pearson’s correlations between them were calculated.

### 2.6 Hub gene selection

We obtained the key genes in the most significant module of the PPI network. The phenotype-related modules in the WGCNA network were also identified, and the genes in those modules were extracted. Hub genes common to both networks were selected as candidates for further analysis and validation.

### 2.7 Coexpression network construction of the TCGA dataset for further validation

To confirm the reliability of the identified DEGs, we analyzed the TCGA-BRCA data by using the same strategy to obtain the TCGA DEGs. We performed a one-step function of WGCNA for TCGA DEG network construction and the detection of consensus modules. In addition, the correlation between ME and phenotype (cancer or normal) was calculated. The candidate genes that also appeared in the TCGA coexpression network were considered to be the true hub genes.

### 2.8 Kaplan-Meier (KM) survival analysis

The online survival analysis software program called Kaplan-Meier plotter (http://kmplot.com/) contains and utilizes expression data from 5,143 breast cancer patients[16]. The median expression level was used to divide patients into two groups, and overall survival analysis was performed to determine the connection between the expression level of hub genes and the overall survival time of patients. The hazard ratio (HR) was provided, and the P-value was calculated by log-rank tests.

## 3. Results

### 3.1 DEG filtering

With thresholds of |logFC| ≥ 2 and Bonferroni adjusted P-value < 0.01, we extracted 540 and 2,509 DEGs from the expression profile datasets GSE10180 and GSE65194, respectively. Scatter volcano plots were developed to illustrate the distribution of each gene on logFC and -log(P-value) values (Figure 1A). After integrated bioinformatics analysis, a total of 322 consistently DEGs were identified from the two datasets (Table S1). Among those DEGs, 69 are up-regulated and 253 are down-regulated. In addition, the gene expression pattern was consistent in both datasets, as shown in the heatmap (Figure 1B, 1C).

**Figure 1.**
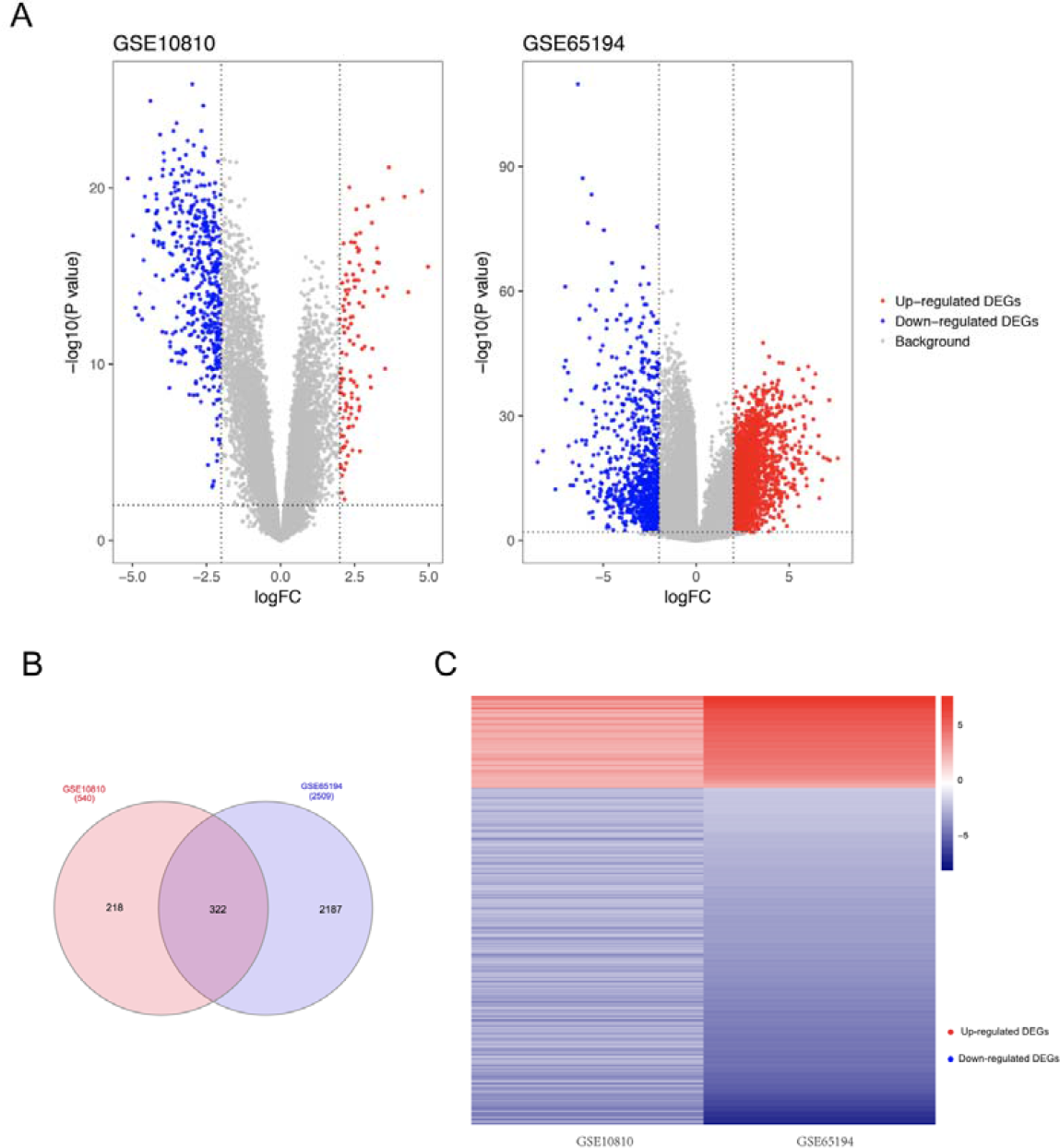
DEGs in each GEO dataset and common DEGs between two GEO datasets. **A.** Volcano plot of DEGs in each GEO dataset**. B.** Common DEGs shared by two datasets. **C.** Common DEGs in both datasets with the same gene expression pattern (red: up-regulated genes, blue: down-regulated genes).

### 3.2 KEGG pathway analysis and GO enrichment analysis

To explore the DEG functions, we performed KEGG pathway and GO enrichment analysis. The top results of each functional group are shown in Figure 2A and Table S2. Oocyte meiosis, cell cycle and progesterone-mediated oocyte maturation are the pathways in which up-regulated genes are mainly enriched. Most of the down-regulated genes were enriched in the PPAR signaling pathway, AMPK signaling pathway, regulation of lipolysis in adipocytes and adipocytokine signaling pathway. The GO enrichment results showed that most of the up-regulated genes were enriched in nuclear division, mitotic nuclear division, organelle fission and the regulation of nuclear division, and the down-regulated genes were mainly enriched in lipid localization.

**Figure 2.**
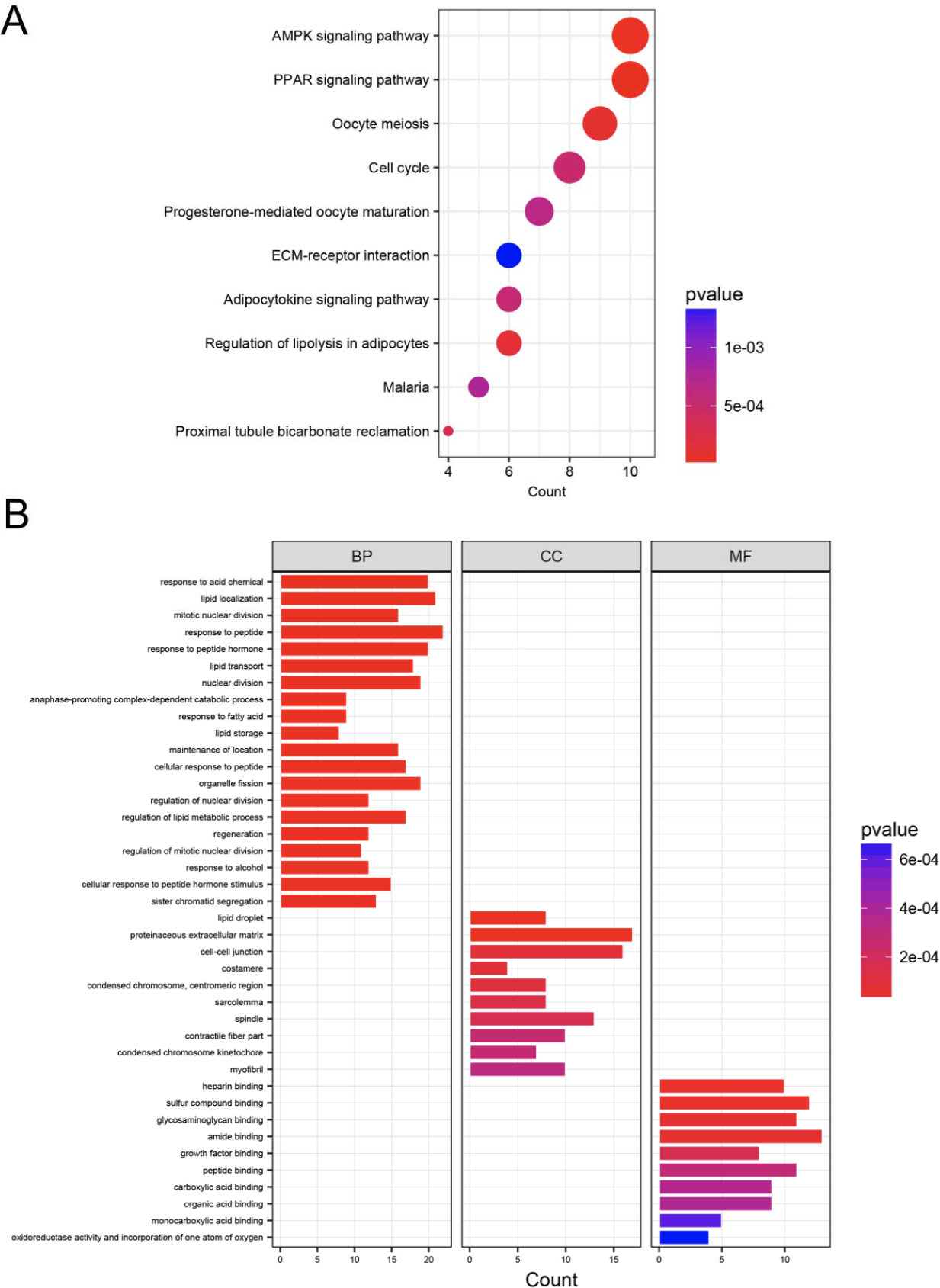
Top KEGG pathway enrichment and GO functional enrichment of 322 common DEGs. **A.** Top KEGG pathway enrichment results of 322 DEGs. **B.** Top enriched GO terms of key DEGs classified into the molecular function (MF) group, the biological process (BP) group and the cellular component (CC) group.

### 3.3 Identification of key DEGs and significant clusters in the PPI network

STRING database was used to obtain the interactive relationships of DEGs. Genes with a combined score > 0.9 were defined as key DEGs. Finally, 95 key DEGs as network nodes and 244 edges were used to construct the PPI network (Figure S1A). MCODE recognized three of the most significant clusters and identified 28 genes = from the PPI network (Figure S1B, Table 1).

**Table 1.**
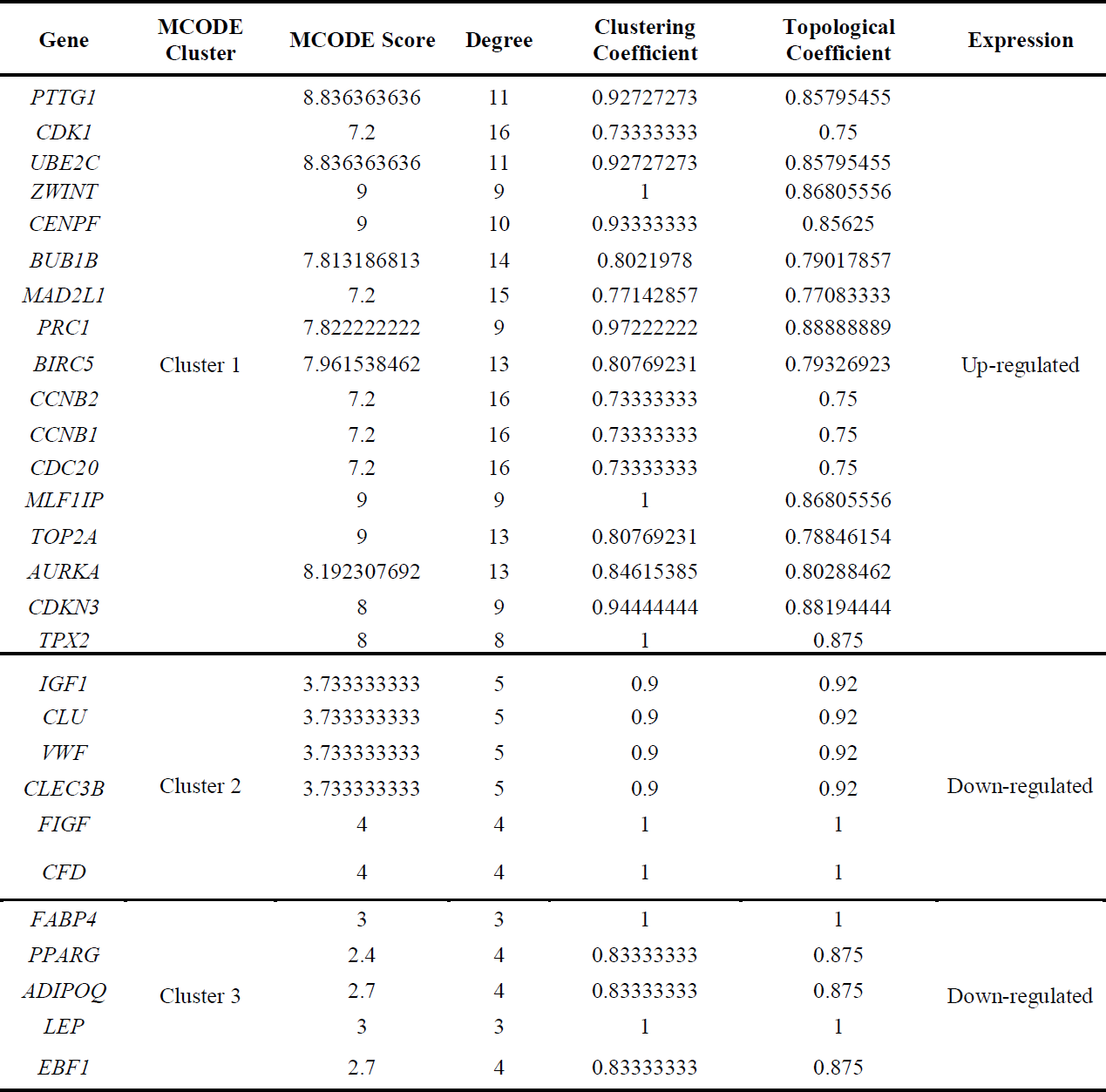
The key genes of DEGs identified from the PPI network.

### 3.4 Construction of the weighted coexpression network and identification of key modules

WGCNA analysis was used to classify the DEGs into different modules based on their similarity of expression patterns by performing the method of average linkage clustering. In this study, three modules were identified according to the breast cancer phenotype (Figure 3A). We considered the blue module of the MEs to be the one we found, due to its highest correlation with breast cancer (Figure 3B). Finally, a total of 35 genes were identified in the blue module (Table 2) and were considered the most relevant genes for breast cancer.

**Table 2.**
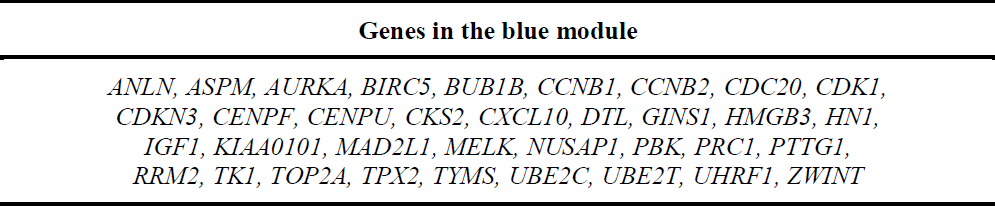
The genes identified from the weighted coexpression network.

**Figure 3.**
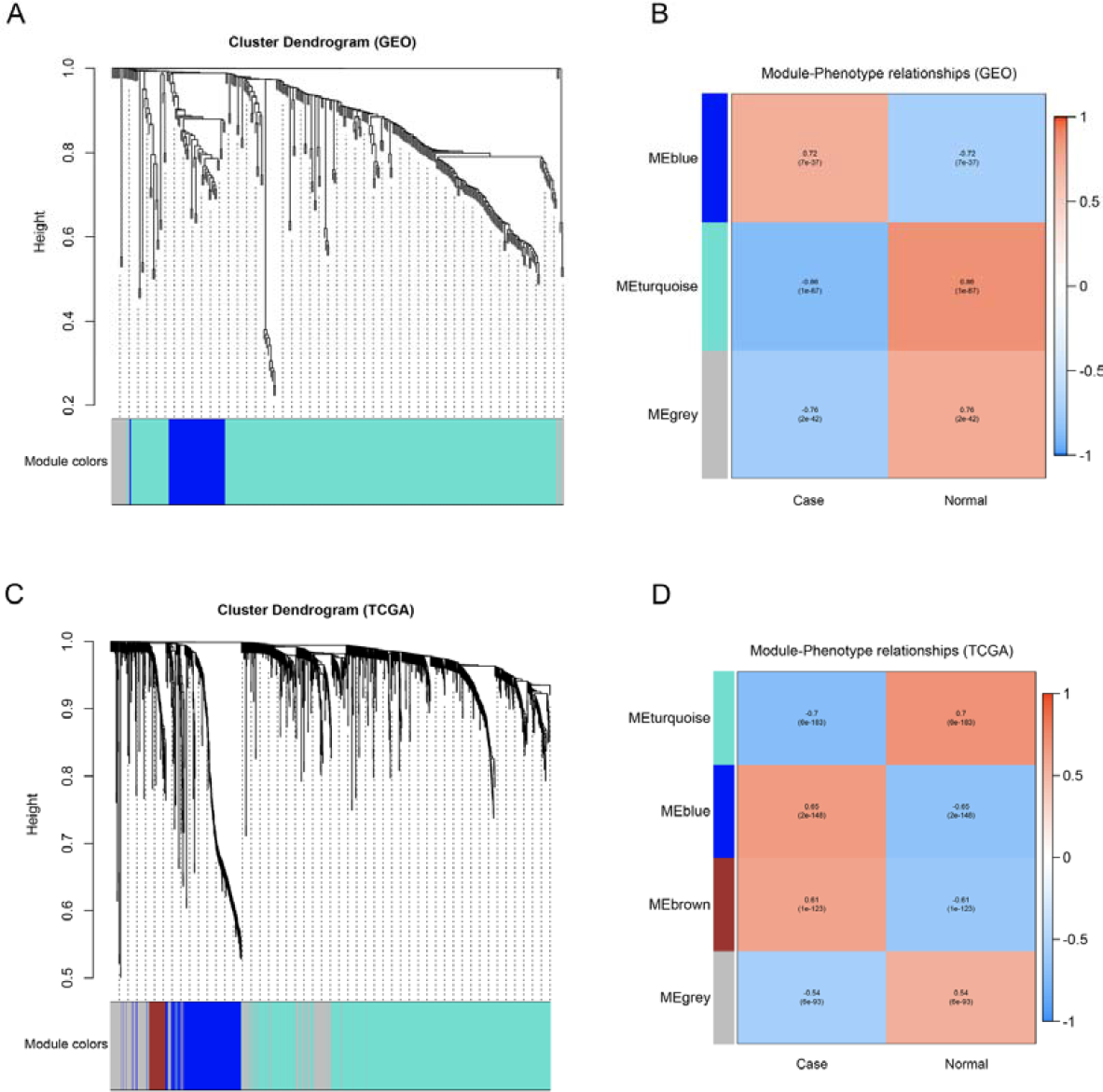
WGCNA coexpression network analysis of the GEO datasets and TCGA dataset. **A.** Gene dendrogram obtained by clustering the DEGs in the GEO datasets. **B.** Relationships of consensus module eigengenes and phenotype in the GEO datasets. **C.** Gene dendrogram obtained by clustering the DEGs in TCGA dataset. **D.** Correlations of consensus module eigengenes and phenotype in TCGA dataset.

### 3.5 Hub gene selection

According to the WGCNA results, a total of 35 genes were considered to be highly correlated with the blue module (Table S3). We then found that 17 genes identified from the PPI network are consistent across the WGCNA network. Hence, 17 common network genes were considered hub genes, subject to further analysis and validation.

### 3.6 Coexpression network construction of the TCGA dataset for further validation

The coexpression network of the differential expression profile data from TCGA was also constructed by WGCNA with the same approach described in this report, and four modules were found to be related to the breast cancer phenotype (Figure 3C). The MEs of the blue and brown modules exhibited a much higher correlation with breast cancer phenotype than the other modules (Figure 3D). Upon integrating the 17 hub genes obtained from the PPI and WGCNA network above, the genes *MAD2L1* and *IGF1* were excluded, and the remaining 15 (*AURKA, BIRC5, BUB1B, CCNB1, CCNB2, CDC20, CDK1, CDKN3, CENPF, PRC1, PTTG1, TOP2A, TPX2, UBE2C* and *ZWINT*) were found in the blue module. Based on both methods, we finally identified 15 hub genes for breast cancer.

### 3.7 Kaplan-Meier (KM) survival analysis

To further evaluate the prognostic importance of the genes that we considered hub genes in this report, overall survival analysis was applied to investigate the association of these genes with the overall survival time of breast cancer patients by using Kaplan-Meier plotter (Figure 4). We found that all the hub genes with higher expression levels were associated with significantly shorter overall survival time among breast cancer patients, which might suggest that these hub genes are closely related to breast cancer.

**Figure 4.**
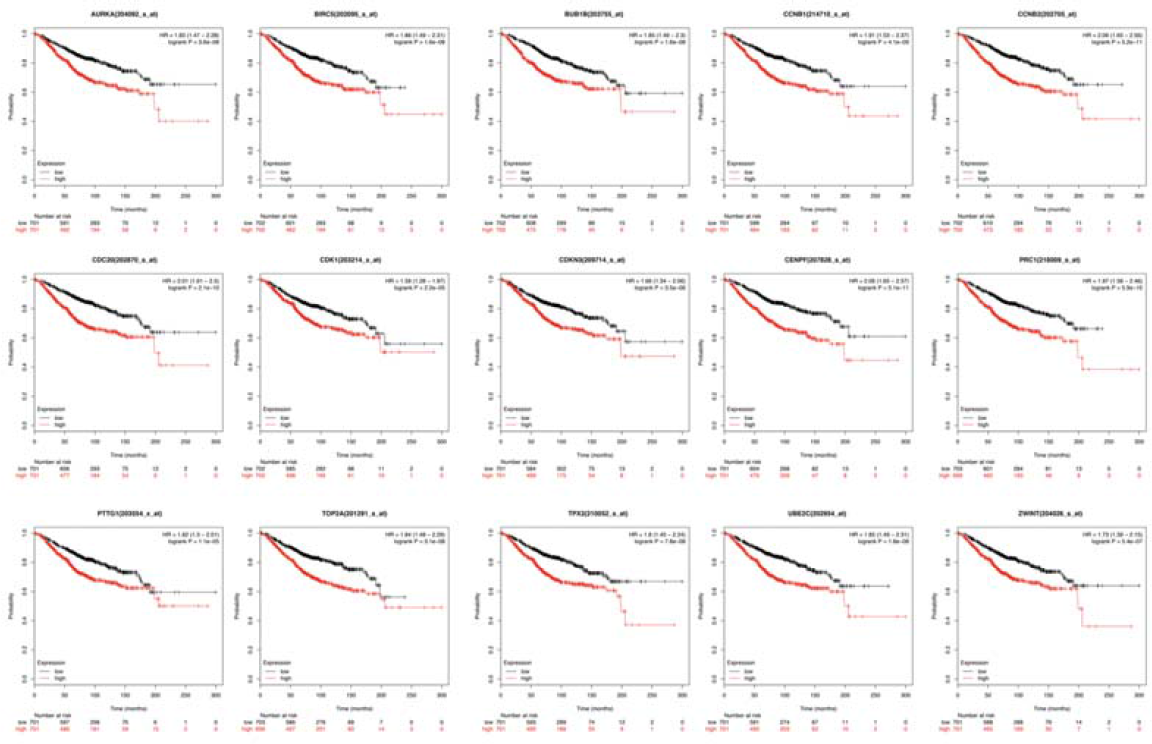
The overall survival of hub genes in breast cancer.

## 4. Discussion

Although the treatment of breast cancer has improved markedly, it remains the most prevalent malignant tumor with the highest increase in prevalence among women worldwide. Uncovering the molecular mechanisms of breast cancer is critical to its diagnosis, therapy and prognosis. The DNA microarray gene expression profile has proven its value and is widely used to explore differentially expressed genes involved in tumorigenesis, which will provide valuable information for clinical applications.

In this study, two gene expression profile datasets (GSE10810 and GSE65194) from the GEO database were retrieved and analyzed. We filtered common DEGs and then identified 17 hub genes that were detected in both the PPI and WGCNA coexpression networks. To further validate these genes, we extracted TCGA-BRCA data to screen the modules related to the phenotype of breast cancer by using WGCNA. After this comparison, 15 real hub genes that were closely associated with breast cancer were identified. This finding may provide valuable information for treatment decisions and prognosis predictions regarding breast cancer.

Notably, these 15 hub genes were all commonly overexpressed among breast cancer patients. KEGG enrichment analysis demonstrated that these hub genes were mostly related to the cell cycle, oocyte meiosis and p53 signaling pathways, and GO enrichment analysis also revealed that they were significantly involved in the cell cycle, cell division, nuclear division and chromosome segregation processes (Figure S2, Table S4). These results suggested that the hub genes were highly related to chromosome instability and probably play an irreplaceable role in tumorigenesis and tumor proliferation. Furthermore, by performing KM survival analysis on these hub genes, we also found that higher expression of those genes was associated with a worse prognosis among breast cancer patients. All of these results indicated that the 15 hub genes might be closely associated with breast cancer and could be potential biomarkers for prognosis.

We also identified that three (*BUB1B, TOP2A* and *AURKA*) of the 15 hub genes were commonly found in the OncoKB cancer gene list[17]. This finding could validate our conclusion from another perspective. *BUB1B* encodes a kinase that is related to the spindle checkpoint function and controls proper chromosome segregation during cell division[18]. The protein encoded by this gene is localized to the kinetochore and is involved in anaphase-promoting complex/cyclosome (*APC/C*) inhibition, which delays the onset of anaphase and ensures proper chromosome segregation. Thus, it plays important roles in tumor proliferation and progression among multiple cancer types[19]. As a checkpoint-related gene, *BUB1B* overexpression might increase the risk of cancer. *TOP2A* controls DNA topologic states and cell progression[20]. This nuclear enzyme is mainly related to processes such as chromatid separation, chromosome condensation, and the relief of torsional stress that occurs during DNA transcription and replication. The upregulation of *TOP2A* was associated with female breast cancer and other cancer types[21]. As a negative regulator of p53, *AURKA* promotes tumor growth and cell survival. *Myc* and *AURKA* regulate each other’s expression at the transcriptional level and contribute to the genesis of liver carcinoma[22].

The remaining 12 hub genes are also important and highly involved in many tumor processes. *PTTG1* prevents separin from promoting sister chromatid separation by encoding securin proteins. *PTTG1* promotes tumor cell growth and malignancy in breast cancer[23]. *CDK1* promotes cell cycle gene expression and is necessary for faithful cell division[24]. Targeting *CDK1* can inhibit the cellular proliferation of liver cancer cells. As a member of the *E2* ubiquitin-conjugating enzyme family, the protein that *UBE2C* encodes is highly involved in mitotic cyclin disassembly and the cell cycle. Hence, *UBE2C* might affect the progression of cancer to some extent. *BIRC5* is a protein-coding gene from the inhibitor of apoptosis (IAP) gene family. It functions as a negative regulator that prevents the cell from undergoing apoptosis [25]. *CCNB1* and *CCNB2* are both members of the cyclin family. As essential components in cell cycle regulation, *CCNB1* and *CCNB2* appear to act as oncogenes and are highly associated with breast cancer according to many studies[26]. Acting as a regulatory protein during cell cycle progression, *CDC20* performs certain functions in coordination with a series of other proteins. Moreover, it has been suggested to affect breast cancer survival. *ZWINT*, however, is believed to be involved in kinetochore function, although the detailed mechanism remains unknown. A study revealed that *ZWINT* overexpression affects cell proliferation in breast cancer[27]. *CENPF* protein is required for kinetochore function during cell division and is related to the cell cycle, mitotic, and cell proliferation pathways. Together with *FOXM1*, these genes become copilots driving cancer malignancy[28]. *PRC1* encodes a protein that is involved in cytokinesis and is essential for cell cleavage. *PRC1* overexpression was detected in p53-defective cells, and a negative regulation cycle was found to be controlled by p53[29]. *CDKN3* encodes a cyclin-dependent kinase inhibitor protein and is essential for normal mitosis and G1/S transition[30]. Its overexpression in human cancer usually indicates poor survival for patients. Therefore, it is also a target in cancer treatment research. *TPX2* is a spindle assembly factor that could serve as a prognostic marker and promote proliferation, progression, migration and invasion in breast cancer.

In conclusion, this study identified 322 consistent candidate DEGs and finally revealed 15 hub genes by using multiple cohort profile datasets and a series of bioinformatics analyses. These hub genes were significantly associated with the cell cycle, oocyte meiosis, and p53 signaling pathways and were also significantly enriched in the cell cycle, cell division, nuclear division, chromosome segregation and other tumor-related processes, which might prove their value in clinical applications involving breast cancer. This finding would effectively promote the understanding of the inner cause of breast cancer, and the 15 hub genes might serve as cancer biomarkers for prediction, diagnosis, individualized prevention, therapy and prognosis.

## Acknowledgements

We would like to thank all the members for their enthusiastic participation in this work. This research was supported by Shenzhen Municipality Government of China (JSGG20140702161347218).

